# First record of *Culex pipiens* (Diptera: Culicidae) in Alberta: Expanding distributions and ecotype patterns in a western Canadian province

**DOI:** 10.1101/2024.09.24.614553

**Authors:** Ty Pan, Michaela Seal, Hailey Shaw, Shahaanaa Mohanaraj, Gen Morinaga, Brittany Hogaboam, Michael Jenkins, Alexandra Coker, John Soghigian

**Affiliations:** Faculty of Veterinary Medicine, University of Calgary, Calgary, Alberta, T2N 1N4; Integrated Pest Management Lab, City of Edmonton, Edmonton, Alberta, T5G 2S7; Parks and Open Spaces, City of Calgary, Calgary, Alberta, T2P 2M5

## Abstract

*Culex pipiens* is an invasive mosquito found in temperate regions globally. It is considered among the most important disease vectors worldwide and is responsible for transmission of a range of pathogens, including West Nile virus, avian malaria, Saint Louis encephalitis, and filarial worms. Throughout its northern temperate range, this mosquito is found in two ecotypes: form *pipiens* and form *molestus*. In Canada, this mosquito was previously thought restricted to the Pacific coast of British Columbia and the eastern provinces of Ontario, Quebec, and the Maritimes. Through routine mosquito surveillance and targeted trapping for *Cx. pipiens*, we detected this mosquito in two Albertan municipalities earlier than suggested by species distribution modelling based on climate change data. We confirmed the identity of putative *Cx. pipiens* specimens using DNA sequencing and found that form *pipiens* was the most common ecotype found in Alberta. Further, we compared the frequency of ecotypes in Alberta to elsewhere in North America and found a general trend of increased form *pipiens* in more northern latitudes, similar to previously reported results. We discuss our findings in context of vector-borne disease activity in Canada, particularly West Nile virus.

## Introduction

Anthropogenic activities and the rise in global temperatures is contributing to the rapid expansion of insect vectors of disease (Fecchio et al. 2021). One such species is *Culex pipiens* (Diptera: Culicidae), the northern house mosquito or common house mosquito. This species is native to Europe, western Asia, and North Africa, but invasive in other temperate regions globally (Haba & McBride, 2022). Its wide invasive distribution and the ability for this mosquito to be a successful vector of many pathogens such as West Nile virus, avian plasmodium, and nematodes (Turell et al. 2001) makes this species one of the most important disease vectors to humans and other animals in the northern hemisphere. It is widely considered responsible for the emergence of West Nile virus in North America in 1999 (Fecchio et al. 2021).

*Culex pipiens* is one member of a species complex which includes *Cx. p. pipiens, Cx. p. pallens, Cx. quinquefasciatus, Cx. restuans, Cx. australicus, and Cx. salinarius* (Harbach 2012). These species are similar morphologically, which can make differential identification from morphology alone difficult, particularly when specimens are damaged by trapping. *Culex pipiens* is further divided into two ecotypes, form *pipiens* and form *molestus*, which we refer to collectively as *Cx. pipiens sensu stricto* following Haba and McBride (2022). The females of these two ecotypes are similar in morphology but vary in ecology and can be differentiated via molecular markers (Becker et al. 2020). Form *pipiens* is primarily active above ground, and feeds on birds (Haba and McBride 2022), while in northern latitudes, form *molestus* typically inhabits underground areas (e.g., sewers and train tunnels) and feeds on mammals, including humans (Haba and McBride 2022). Ecotypes appear to be fully reproductively compatible and can hybridize with other closely related species *such* as *Cx. quinquefasciatus* (Haba and McBride 2022). It is very likely that hybrids between ecotypes, and between closely related species, are more efficient vectors of disease, and hybrids have even been associated with West Nile transmission in North America and Europe (Andreadis 2012; Turell 2012).

*Culex pipiens s*.*s*. has historically been restricted in Canada to southern Ontario, the Maritimes, and coastal British Columbia. However, species distribution models informed by climate change predictions indicated that by 2100 all southern Canada is expected to be ideal habitat for *Cx. pipiens s. s*. (Hongoh et al. 2012). Moreover, recent habitat distribution models suggest suitable habitat exists in other provinces for this species (Gorris et al. 2021). *Culex pipiens* utilizes urban environments well, which will likely enable rapid expansion of this species as human development and climate change continue to alter the environment in Canada (Peach and Matthews 2022). Beyond its ability to utilize urban habitats, this species frequently oviposits in smaller temporary water sources, such as flowerpots, bird baths, and various other manmade containers (Haba and McBride 2022). This behavior exposes it to opportunities for human-aided dispersal, and thus further increasing its potential for range expansion. Here, we document the first confirmations of *Cx. pipiens s*.*s*. in the province of Alberta through an initial detection of the species in Edmonton in 2018, and a subsequent detection in Calgary in 2022 through routine and targeted surveillance. We demonstrate via molecular methods that the populations found in this province are a mix of ecotypes and contextualize this within geospatial variation in ecotypes frequencies across latitudes in North America.

## Materials & Methods

Due to the unexpected detection of *Culex pipiens*, our surveillance efforts were reactive and not targeted towards this species. The first detection of *Culex pipiens* specimen was during routine surveillance by the City of Edmonton in 2018. *Culex* mosquitoes that were abundant and looked markedly different from the native *Cx. tarsalis* were set aside by mosquito surveillance technicians. These mosquitoes were identified morphologically as either *Cx. pipiens* or *Cx. restuans*. The finding of *Cx. pipiens* in Edmonton prompted collaboration with the City of Calgary to inspect specimens collected through their routine mosquito surveillance program for *Cx. pipiens*. We identified both adult and larval *Cx. pipiens* from traps in Calgary in the summer of 2022 using morphological identification (Darsie and Ward, 2005). We confirmed the morphological identifications of *Cx. pipiens* in Edmonton and Calgary with routine barcoding of the cytochrome oxidase subunit 1 marker (Folmer et al. 1994), which we sequenced at the University of Calgary’s Centre for Health Genomics. We assembled the resulting sequencing reads in Geneious Prime (Geneious Prime 2023.0.4) and compared then to the NCBI nucleotide database with BLAST (Altschul et al., 1990). In addition, we used molecular ‘ecotyping’ methods as described in Bahnck and Fonseca (2006) to characterize the ecotype of *Cx. pipiens* found in Alberta.

To compare the frequency of *Cx. pipiens* ecotypes we found in Alberta to elsewhere in North America, we conducted a literature review of *Cx. pipiens* ecotypes across North America. We searched Google Scholar with the keywords “North America” and “CQ11”, “Canada” and “CQ11”, and “United States” and “CQ11”. We only selected studies that included *Culex pipiens* specimens collected in North America and molecularly analyzed using CQ11 primers. Nine papers met our criteria and provided enough information for analysis (Supplemental Table 1). We extracted the sampling location, microhabitat (i.e. above or below ground), and the proportion of form *pipiens* and form *molestus* specimens from each article. We also included data from the molecular analysis of Alberta specimens from Calgary and Edmonton in the analysis. We calculated the frequency of each ecotype for each location, and geospatial analyses of the data were completed in R version 4.3.2 using *ggplot2* (v3.4.4; Wickham 2024) and *Leaflet* (v2.2.1; Cheng et al. 2024) packages.

## Results & Discussion

We confirmed the identity of *Culex* mosquitoes collected in Edmonton in 2018 as *Cx. pipiens* (Supplemental Table 2). In Calgary, we first detected this mosquito in 2022 and confirmed via barcoding (Supplemental Table 2). Since its first detection in both municipalities, the mosquito has become widespread in both municipalities and is commonly found during surveillance activities.

Unfortunately, as our sampling was primarily reactive, we are unable to judge exactly how fast the mosquito has expanded throughout both cities.

Using the CQ11 primers, we confirmed that both ecotypes, form *pipiens* and form *molestus*, are present in Edmonton and Calgary (Table 1). Ecotype *pipiens* was predominant in both cities, however a small percentage of specimens were *molestus*. Calgary and Edmonton had similar hybrid rates at about 15%. Alberta populations were predominantly form *pipiens*, which is consistent with other above-ground populations at more northerly latitudes (Figure 1, also see Supplementary Table 1). This reflects the general trend of increasing form *pipiens* allele frequency in above-ground populations as latitude increases, a phenomenon also observed in European populations (Haba and McBride 2022).

**Table 1.** Identification of Culex pipiens ecotypes using CQ11 primers.

| City | Collection Year | N total | % *pipiens* | % *molestus* | % hybrid |
| --- | --- | --- | --- | --- | --- |
| Calgary | 2022 | 30 | 80 | 6.7 | 13.3 |
| Edmonton | 2018, 2020, 2021 | 173 | 80.3 | 3.5 | 16.2 |

**Figure 1.**
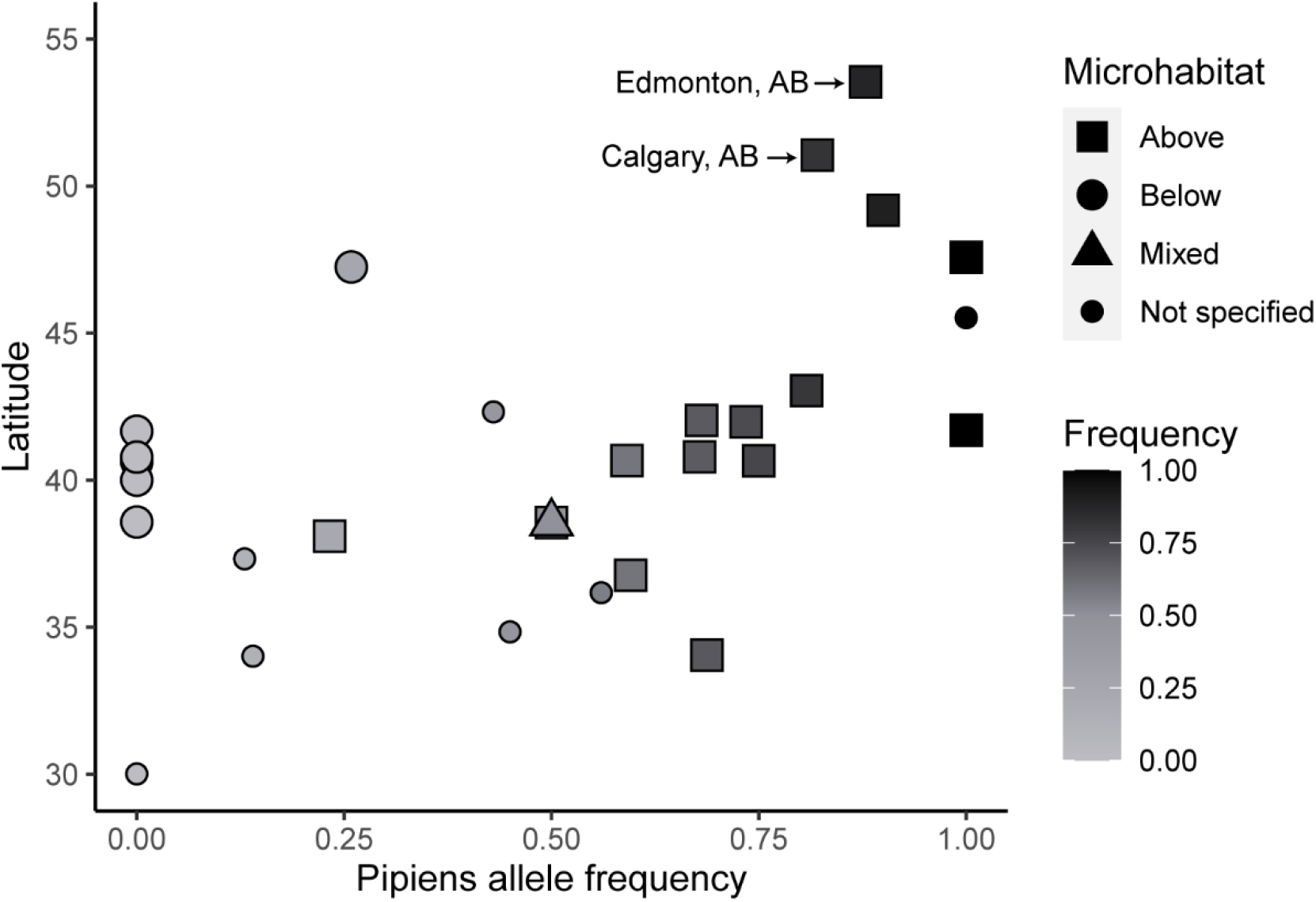
Frequency of *pipiens* alleles across latitude and microhabitat for *Culex pipiens* found in North America based on analysis with CQ11 primers from 2006 to 2022. *Culex pipiens* sampled from Calgary and Edmonton, Alberta, Canada, predominantly had the *pipiens* allele, but with a notable percentage of *molestus* alleles and hybrids (see Table 1).

Our finding of a small percentage of *Cx. pipiens* as form *molestus* in Alberta is surprising, as neither city has significant underground subway systems, which are commonly associated with this form. However, both cities have light rail systems that travel briefly underground, as well as substantial numbers of underground parking garages, which may serve as habitat for these mosquitoes. Interestingly, we captured all our mosquitoes above-ground, which indicates that form *molestus* in Alberta is not confined to underground localities. This has been observed in some locations in Europe as well, albeit at a lower latitude (Haba and McBride 2022). Alternatively, admixture among *pipiens* and *molestus* populations elsewhere in North America, and subsequent gene flow, may have made the CQ11 locus less informative for ecotype discrimination; further genomic analyses will be necessary to evaluate this possibility.

As global temperatures rise, vector-borne diseases pose an increased risk as mosquitos such as *Culex pipiens* expand their range (Fecchio et al. 2021). Additionally, increasing global temperatures will promote development of mosquito larvae (Becker et al. 2020) and may increase feeding rates and reproduction of adult vectors (Mordecai et al. 2017). Owed to its vector competence for West Nile virus, *Cx. pipiens* is of particular concern as climate change is expected to triple West Nile virus cases in the continental United States over the next 30 years (Paull et al. 2017). West Nile virus was first detected in Canada in 2001 with particularly high cases in 2003 and 2007 (Giordano et al. 2017; Public Health Agency of Canada 2024). West Nile virus now causes only sporadic human and horse cases in Canada, and as a result, surveillance for both the vectors and the pathogen has declined throughout much of Canada. In light of previously reported expansion of invasive mosquito species in Canada (Peach and Matthews 2022), and the expansion we report here for *Cx. pipiens*, a review and potential expansion of vector and virus surveillance may be warranted in Canada generally. Furthermore, *Cx. pipiens* is an efficient vector for avian malaria. Although typically thought to be largely harmless to native birds in North America, a new virulent strain of avian malaria has recently been detected in the United States (Theodosopoulos et al., 2021). In addition, exotic birds are at risk – for instance, there have already been deaths recorded of two zoo penguins in Calgary (Wilder Institute/Calgary Zoo 2023), leading to increased concern of the risks *Cx. pipiens* poses to exotic and native birds in Alberta.

Our finding of *Culex pipiens* in Alberta warrants future study, including the potential implications for veterinary and public health, as well as the routes of invasion that this mosquito utilized to arrive in Alberta. We find the timeline of Cx. pipiens particularly intriguing — our findings indicate that *Cx. pipiens* was likely present in Edmonton before Calgary, rendering a northward expansion of a southern population unlikely. One possibility is this mosquito was introduced to Edmonton via an anthropogenic means; alternatively, it may have expanded naturally eastward from British Columbia. The introduction in Calgary could be due to range expansion from the Edmonton population, anthropogenic movement, or a combination of multiple causes. Future studies will investigate the origin of the introduction of *Cx. pipiens* in Alberta through genetic analysis, as well as characterize the ecology of *Cx. pipiens* in an Albertan context.

## Supplementary Material

**Supplementary Figure 1.**
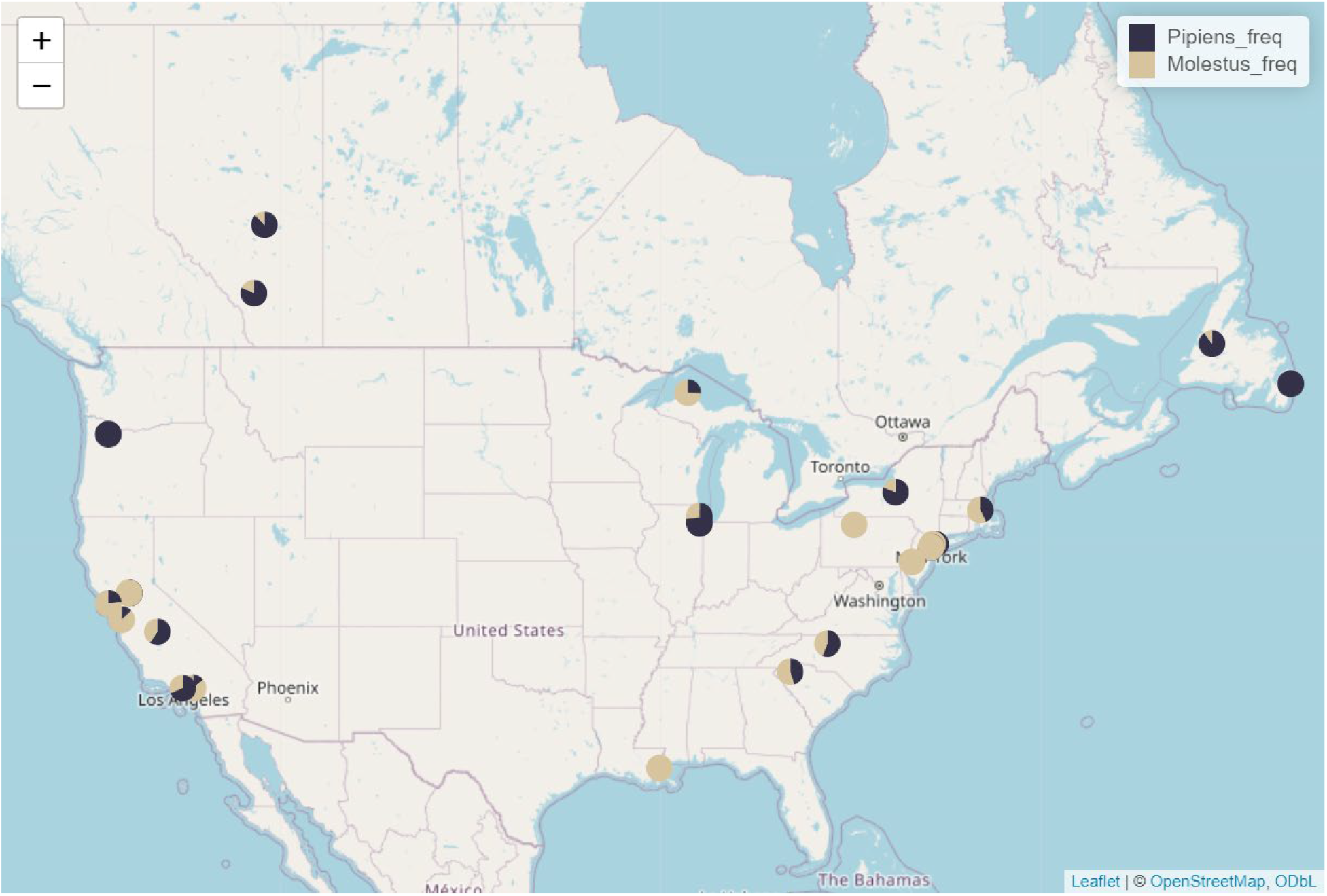
Map of the proportion of ecotypes *pipiens* and *molestus* from *Culex pipiens* populations across North American using analysis with CQ11 primers from 2006 to 2022, including analysis of recently discovered populations of *Culex pipiens* in Alberta.

**Supplementary Table 1.** List of sources used for the analysis of ecotype frequencies across North America and the sampling location used in each source. Only sources that used CQ11 primers for ecotype analysis were used, and papers found were published between 2006 and 2016.

| Source | Sampling location |
| --- | --- |
| Kent, R.J., Harrinton, L.C. & Norris, D.E. (2007) | New York |
| Cornel, A., Lee, Y., Fryxell, R.T., Siefert, S., Nieman, C., Lanzaro, G. (2012) | California |
| Bahnck, C.M. & Fonseca, D.M. (2006) | Massachusetts, California, South Carolina, Louisiana, Pennsylvania, Oregon, North Carolina |
| Chaulk, A.C., Carson, K.P., Whitney, H.G., Fonseca, D.M., & Chapman, T.W. (2016) | Newfoundland |
| Asgharian, H., Chang, P.L., Lysenkov, S., Scobeyeva, V.A., Reisen, W.K., & Nuzhdin, S.V. (2015) | Sacramento, CA |
| Fritz, M.L., Walker, E.D., Miller, J.R., Severson, D.W., & Dworkin, I. (2015) | Illinois |
| Kothera, L., Godsey, M., Mutebi, J., & Savage, H. (2010) | New York |

**Supplementary Table 2.** A subsample of sequenced mosquito specimens with their first top BLAST results. Specimens were collected from Edmonton and Calgary from 2018 and 2022 respectively.

| Lab Code | City | Year Collected | Top BLAST Result | Percent Identity | Accession |
| --- | --- | --- | --- | --- | --- |
| 047ED | Edmonton | 2018 | *Culex pipiens* | 100% | MN005046.1 |
| 049ED | Edmonton | 2018 | *Culex pipiens* | 100% | MN005046.1 |
| 050ED | Edmonton | 2018 | *Culex pipiens* | 100% | MN005046.1 |
| 010CA | Calgary | 2022 | *Culex pipiens* | 100% | MN005046.1 |
| 011CA | Calgary | 2022 | *Culex pipiens* | 100% | MN005046.1 |
| 020CA | Calgary | 2022 | *Culex pipiens* | 100% | MN005046.1 |

## Notes

### Competing Interest Statement

The authors have declared no competing interest.

